# Modeling reveals metabolic basis of competition among *Dehalobacter* strains during tandem CF and DCM metabolism

**DOI:** 10.1101/2025.03.03.641252

**Authors:** Olivia Bulka, Elizabeth A. Edwards, Radhakrishnan Mahadevan

**Author notes:** Corresponding Author: Radhakrishnan Mahadevan.

## Abstract

SC05-UT is an anaerobic mixed microbial enrichment culture that reduces chloroform (CF) to dichloromethane (DCM) through reductive dechlorination, which it further mineralizes to carbon dioxide. This DCM mineralization yields electron equivalents that are used to reduce CF without addition of exogenous electron donor. By studying this self-feeding CF-amended culture and a DCM-amended enrichment culture (DCME), we previously found the genomic potential to perform both biodegradation steps in two distinct *Dehalobacter* strains: *Dehalobacter restrictus* SAD and *Candidatus* Dehalobacter alkaniphilus DAD. Though present in each enrichment culture, strain SAD is more abundant in CF-fed subculture SC05-UT, while strain DAD is more prominent in the DCM-fed subculture DCME. To understand if genomic differences between strains and impact their metabolic mechanisms, the genome of each strain was curated to reconstruct genome-scale metabolic models of each strain, which were then constrained based on thermodynamic and experimental conditions. We demonstrate that metabolic differences between the two strains may allow *Dehalobacter* strain DAD to outcompete strain SAD in the absence of CF, while strain SAD exhibits an advantage in the presence of CF. Additionally, we predict electron cycling methods to reconcile redox imbalances in the cell is required for tandem CF and DCM dechlorination. This work highlights the importance of hydrogen and amino acid exchange in these microbial communities and contributes to the growing body of work surrounding organohalide syntrophy.

**Importance:** Chloroform and dichloromethane contaminate groundwater around the world but can be remediated by microbes capable of metabolizing these toxic compounds. Here we study two distinct strains of *Dehalobacter* and show that while both strains can degrade both chloroform and dichloromethane, differences in their genetic makeup allow each strain to thrive under different environmental conditions. This has implications for understanding the fate of halogenated methanes in the environment and the application of *Dehalobacter* for bioremediation of chlorinated compounds.

## Introduction

The bioremediation industry has long harnessed the power of natural communities for dechlorination of organohalides. Many of these communities contain *Dehalobacter* spp., a key genus for reductive dechlorination of chloroalkanes and alkenes. Despite experimental characterization of more than 40 strains of *Dehalobacter*—and identification of many more strains by 16S amplicon sequencing alone—only six closed genomes have been published within this genus (1–5). Recent pangenomic and phylogenetic work has divided the *Dehalobacter* genus into multiple species: *Dehalobacter restrictus*, *Candidatus* Dehalobacter alkaniphilus, and *Candidatus* Dehalobacter aromaticus (5). *Dehalobacter* are known to be fastidious anaerobic organisms, with fragmented TCA cycles and a reliance on exogenous amino acids and other organic acids (6, 7). Only *Dehalobacter restrictus* has been successfully isolated and deposited in a culture collection, further complicating experimental and computational study of their metabolisms (7–10).

Two genome-scale metabolic models representing organohalide respiring bacterial genera have been reconstructed: a *Dehalococcoides* pangenome model (*i*AI549 (11)), and a *Dehalobacter* pangenome model (*i*HH623 (12)). The *Dehalobacter* model was initially created from the genome of *Ca.* D. alkaniphilus CF and curated to encompass the metabolism of the genus as a whole using annotations from the available genomes of strains CF, DCA, PER-K23, E1, and UNSWDHB (12, 13). Though it has been widely accepted that *Dehalobacter* strains have a very restricted metabolism—each reducing a chlorinated electron acceptor with H_2_ as an electron donor—recent discoveries within the genomes of two novel *Dehalobacter* strains from a mixed microbial culture (SC05) suggest more metabolic variability across species than was previously thought (14).

SC05 is a mixed-microbial anaerobic chloroform (CF)-degrading enrichment culture originally sampled in 2010 from a site polluted with chlorinated ethenes and ethanes (15, 16). Through reductive dechlorination, it reduces CF to dichloromethane (DCM), which it further mineralizes to carbon dioxide and hydrogen (16). This hydrogen can then act as an electron donor to support CF reduction through “self-feeding”, allowing a subculture (SC05-UT) to continually dechlorinate CF to DCM without feeding any exogenous electron donor since 2018 (15). An additional subculture named DCME was established with DCM alone since 2019 (15). Though both of these heterogenous subcultures consist of many microbial species, only a single *Dehalobacter* ASV was detected through 16S amplicon sequencing since their conception; it flourished both during CF dechlorination and DCM mineralization, which nominated it as the sole dechlorinator in the SC05 culture (15).

When the first metagenomes of the two SC05 subcultures (SC05-UT and DCME) were assembled, one *Dehalobacter* metagenome-assembled genome (MAG) emerged from each metagenome (14). Each MAG was searched genomically and proteomically for the enzymes known to perform CF dechlorination—reductive dehalogenases (RDases)—and DCM mineralization—methyltransferases encoded by the DCM catabolism (*mec*) cassette. Only one RDase was expressed in each culture; it was named AcdA in SC05-UT and was shown to dechlorinate CF to DCM (14). The *mec* cassette was also highly expressed in both cultures (14), becoming one of only two reports to ascribe DCM metabolism to *Dehalobacter* (15, 17).

Through additional sequencing, these MAGs were curated into two distinct closed genomes belonging to different *Dehalobacter* species: *D. restrictus* SAD and *Ca.* D. alkaniphilus DAD (5, 18). Both genomes share the RDase–*mec* cassette gene neighbourhood—a region with 98.7% sequence identity across *Dehalobacter* species—in an integration “hot spot” detected in the *Dehalobacter* pangenome (5). Though encoding nearly identical functional genes, these strains each dominate the culture under different conditions—strain SAD is abundant in CF-fed SC05-UT, while strain DAD is more prominent in the DCM-fed DCME (5). Despite this differential predominance, their mutual presence in both cultures challenges the assumption that only one strain is actively growing in each subculture (5).

Here, we aim to quantify these *Dehalobacter* strains in each subculture and rationalize their varying predominance using experimental data coupled with metabolic models, which can be optimized to predict the growth potential of each *Dehalobacter* strain under varying experimental conditions in SC05-UT and DCME. Furthermore, we seek to identify differences in metabolic potential across these two *Dehalobacter* species, to determine ecological significance.

## Results

### Both strains of *Dehalobacter* perform two remediation steps

Strain-specific growth was measured during each remediation step in replicate SC05-UT and DCME subcultures. All bottles were initially fed CF and hydrogen, and *Dehalobacter* growth was quantified during the sequential degradation of CF and DCM using qPCR to target a divergent single copy core gene (*flgC* (15)). Each culture can grow under three defined growth modes (Figure 1A)—Mode 1: supplied H_2_ as an electron donor, CF as an electron acceptor; Mode 2: supplied DCM as electron donor, which provides electrons to reduce CF as the ultimate electron acceptor; Mode 3: supplied DCM as electron donor for mineralization to CO_2_ with H^+^ as the electron acceptor. Under normal maintenance, DCME usually operates in Mode 3, as it is typically only fed DCM and SCO5-UT usually operates in Mode 2 (Figure 1A).

**Figure 1.**
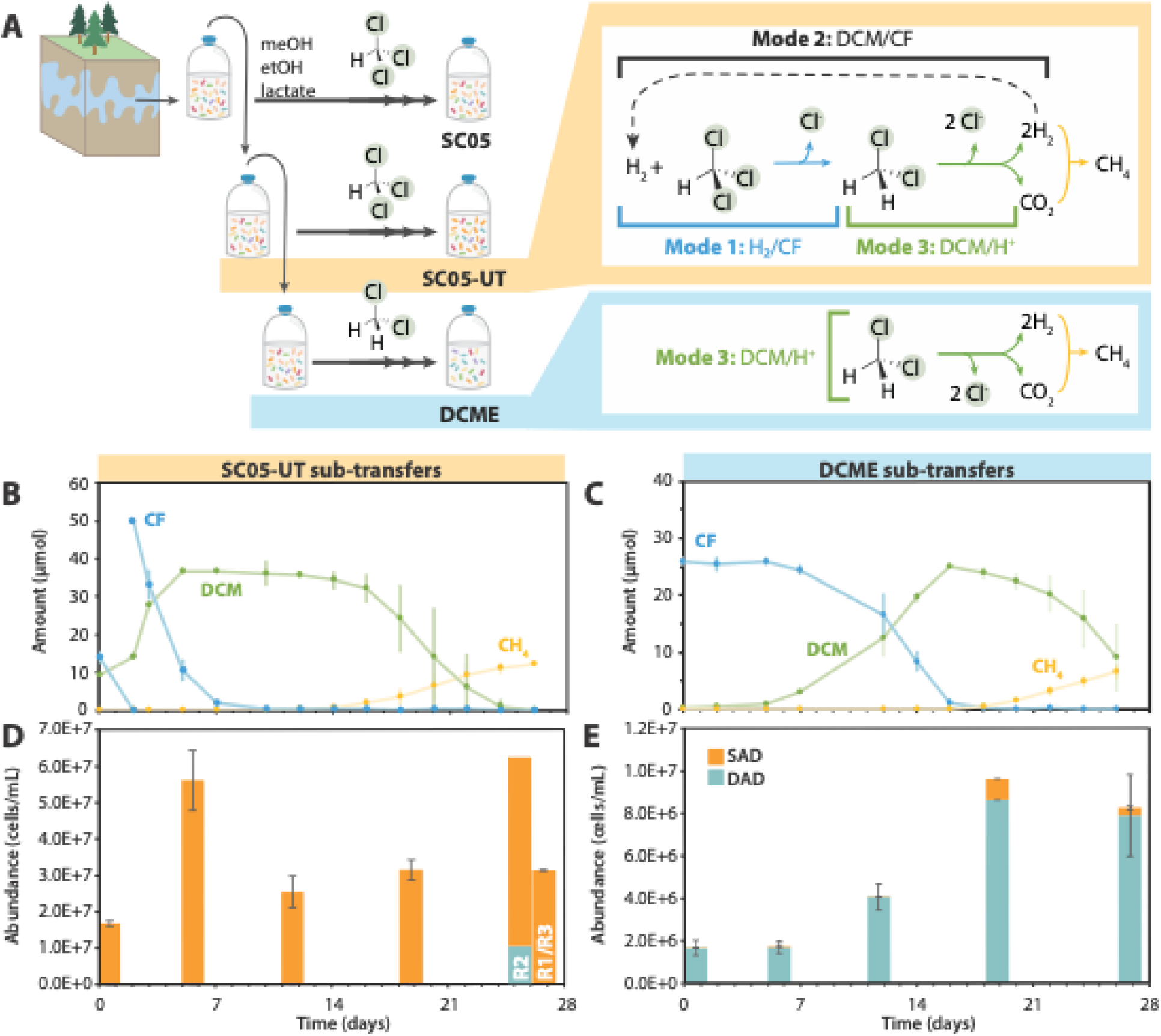
Growth of *Dehalobacter* in SC05-UT and DCME, including **A)** a schematic of culture provenance and metabolic modes, **B)** dechlorination in SC05-UT sub-transfers and **C)** DCME sub- transfers. Panels **D** and **E** show the *Dehalobacter* population in SC05-UT and DCME, respectively. SC05-UT: *n =* 3, error bars represent standard deviation; DCME: *n =* 2, error bars represent range. Day 26 samples from SC05-UT are separated by replicate (R1-R3) to showcase two outcomes.

In the three SC05-UT bottles, only strain SAD grew during CF dechlorination (Figure 1D). The measured cell yield (Table 1) is within error of previously quantified yields of *Dehalobacter* in SC05 when growing on CF alone, and 4-fold to 10-fold higher than that of other CF-dechlorinating strains of *Dehalobacter* (15, 19–22). Some DCM produced from CF dechlorination was consumed concurrently, which may account for the higher yield (Figure 1B). Strain DAD did not grow during this period, which shows unequivocally that strain SAD is the key CF dechlorinator in SC05-UT. During DCM mineralization (Mode 3), strain DAD did not grow in two of three replicates, but in one replicate (R2), strain DAD increased almost 100-fold (Figure 1D, Table 1). Although demonstrating that both strains are independently able to grow by DCM mineralization, this outcome diversity suggests that DCM mineralization may at times be a joint effort between the two *Dehalobacter* strains in SC05-UT.

**Table 1.**
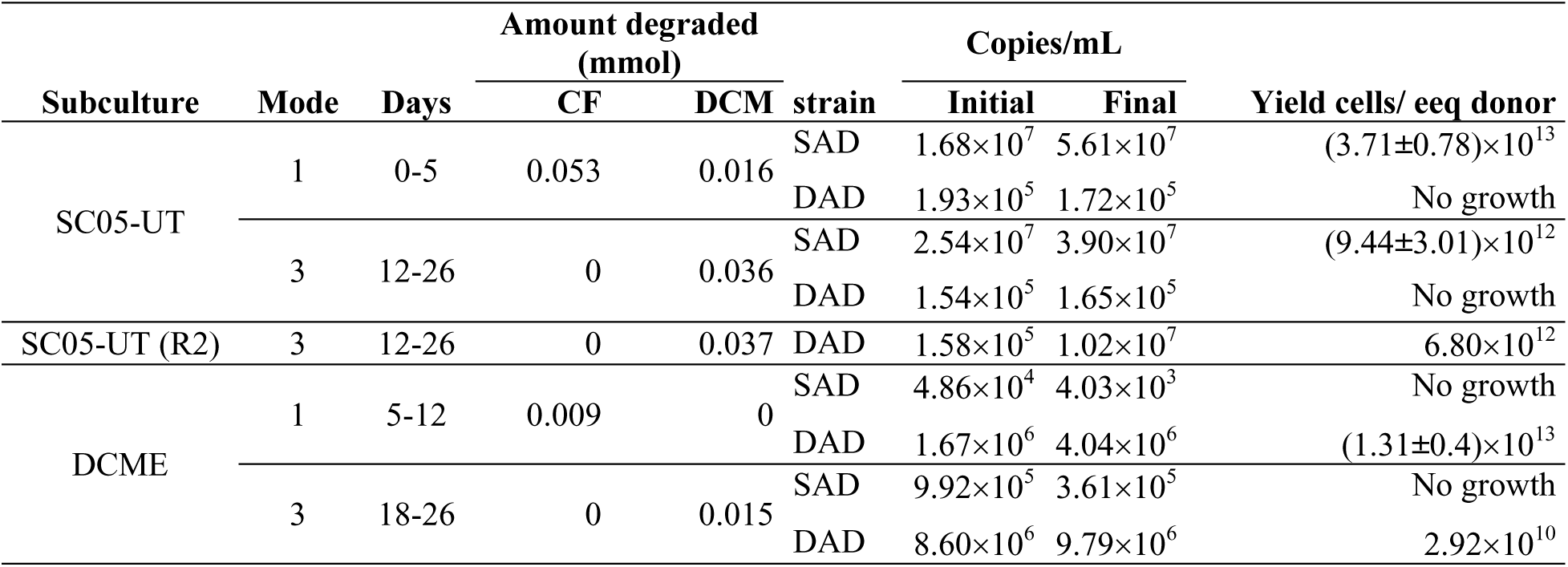
Calculated yields of each strain in SC05-UT and DCME subcultures. Yield is displayed as mean ± error of biological replicates (DCME, *n =* 2 [range]; SC05-UT, *n =* 3 [standard deviation]; no error is shown where one replicate is available).

In the DCME sub-transfers, *Dehalobacter* strain DAD more than doubled as CF was dechlorinated from day 7 to 12 (Figure 1E, Table 1). No DCM was degraded at this time, indicating that strain DAD grows while dechlorinating CF using H_2_ as a donor (Mode 1). During DCM mineralization, strain DAD increased slightly (Table 1), though this yield is lower than previously calculated yields ([1.23 ± 0.65] × 10^13^ cells/eeq donor (15)). Notably, its yield in SC05-UT R2 is cohesive with previous work (Table 1). Though *Dehalobacter* strain SAD was initially 100-fold less abundant than strain DAD in the DCME sub-transfers, strain SAD increased as CF was dechlorinated from day 12 to 19 (Figure 1E, orange). Because strain DAD also grew during this period, its yield cannot be disentangled. Nonetheless, strain SAD’s rapid growth despite low relative abundance suggests it may dechlorinate CF faster or more efficiently than strain DAD, thus outcompeting it when SC05-UT is maintained on CF alone.

### Distinct *Dehalobacter* species metabolic models constructed for each strain

Species-specific genome-scale metabolic models were reconstructed to describe *Dehalobacter* strains SAD and DAD (Table 2) and curated through comparative genomics with support from proteomics (described in Supplemental Texts S1 and S2, respectively). Organohalide metabolism of each *Dehalobacter* strain was updated; reductive dechlorination of CF to DCM and DCM assimilation to methylene-THF are each represented by one reaction (Table S4). A reaction was added to represent a ferredoxin-mediated complex I-like enzyme, reflecting recent work describing electron transfer in *Desulfitobacterium* (a closely related genus) and *Dehalobacter restrictus* (23). Both strain’s genomes encoded a complete Wood-Ljungdahl Pathway (WLP) and 8 homologous hydrogenases (Table S5). These hydrogenases—labelled Hyd-1 through Hyd-8—comprise 4 metabolic reactions (Table S6, Figure S2), as classified by HYDdb (24). One additional hydrogenase was detected in strain DAD (Hyd-9, Table S6), but its representative reaction is redundant. Four hydrogenases were expressed in each of the strains’ proteomes (Table S6). Genes encoding three metabolic reactions were detected uniquely in the genome of strain DAD and were added to the model accordingly: fumarate reductase, aspartate ammonia lyase, and a nitrogen fixation cassette (Figure 2, Figure S1). No unique metabolic genes were found in strain SAD.

**Figure 2.**
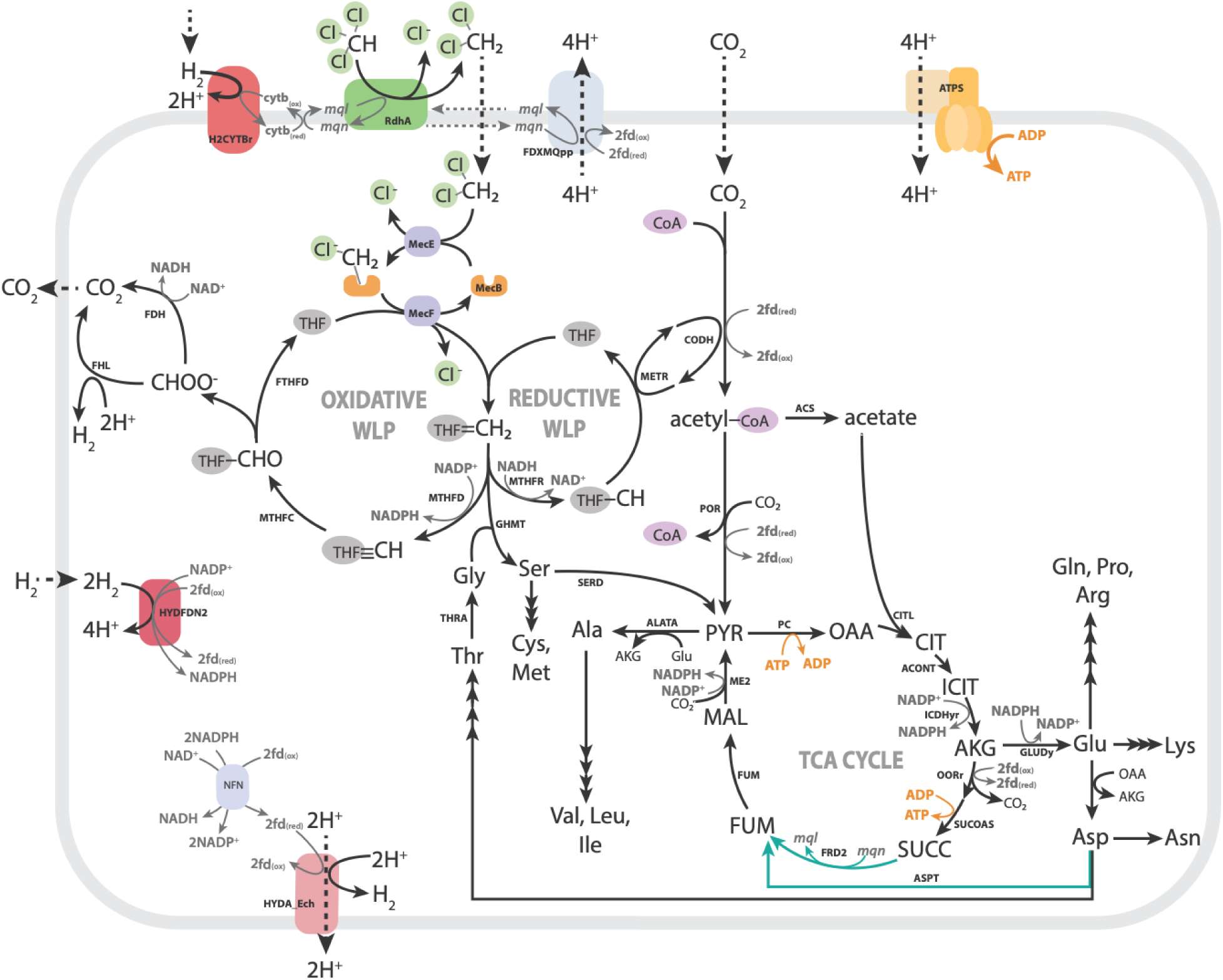
Genetic landscape of central carbon and amino acid metabolism in *Dehalobacter* strains SAD and DAD. Blue pathways are encoded by strain DAD only (ASPT and FRD2). Metabolites and enzymes are abbreviated as follows: PYR: pyruvate, OAA: oxaloacetate, CIT: citrate, ICIT: isocitrate, AKG: alpha-ketoglutarate, SUCC: succinate, FUM: fumarate, MAL: malate, THF: tetrahydrofolate, MecEBF: *mec* cassette, MTHFD: methylene-THF dehydrogenase, MTHFC: methenyl-THF cyclohydrolase, FTHFD: formyl-TFH deformylase, FHL: formate hydrogen lyase, FDH: formate dehydrogenase, MTHFR: methylene-THF reductase, METR: 5-methyl-THF corrinoid/ FeS protein methyltransferase, CODH: carbon monoxide dehydrogenase/ acetyl-CoA synthase, ACS: acetate-CoA ligase, POR: pyruvate oxidoreductase, PC: pyruvate carboxylase, CITL: citrate lyase, ACONT: aconitrate hydratase, ICDHyr: isocitrate dehydrogenase, OORr: oxoglutarate oxidoreductase, SUCOAS: succinate CoA ligase, FRD2: fumarate reductase, FUM: fumarate hydratase, ME2: malic enzyme, ALATA: alanine:oxoglutarate aminotransferase, GLUDy: glutamate dehydrogenase, ASPT: aspartate ammonia lyase, THRA: threonine acetaldehyde lyase, GHMT: methylene-THF:glycine hydroxymethyltransferase, SERD: serine ammonia lyase, NFN: electron-bifurcating transhydrogenase, HYDA_Ech/HYDFDN2/H2CYTBr/HYD-NADH: hydrogenases, FDXMQpp: complex I-like enzyme, RDase: reductive dehalogenase, ATPS: ATP synthase. WLP: Wood-Ljungdahl Pathway, TCA: Tricarboxylic acid.

**Table 2.** Statistics for each curated genome-scale model.

|  | *Dehalobacter restrictus* SAD<br>iOB638 | *Ca. Dehalobacter alkaniphilus* DAD<br>iOB649 |
| --- | --- | --- |
| Genes | 638 | 649 |
| Total reactions | 1068 | 1071 |
| Exchange reactions | 63 | 63 |
| Compartments | 3 | 3 |
| Metabolites | 952 | 952 |

### DCM is a less efficient electron donor than hydrogen

Because these strains grow under three different electron donor/acceptor modes, each model was curated using the thermodynamic constraints for each mode individually (Table **3**; expanded upon in Supplemental Text S1). Energy transfer efficiency is calculated using ATP/e- ratios, which represent the amount of ATP produced per mol electrons transferred from the electron donor (calculated by [model-predicted ATP/*e^-^*]/[max ATP/*e^-^*]). Energy transfer efficiency is lower during Modes 2 and 3—where DCM assimilation occurs—than during Mode 1 (Table **3**). Thermodynamically, DCM mineralization has a favorable **Δ**G, but the cell’s mechanism of metabolism may be unable to harness this to its full extent. Computationally, the model is restricted to using the same reactions as in Modes 1 and 3, thus cannot harness all energy theoretically available. Physiologically, this may be due to these same enzyme mechanism limitations, but we lack clear experimental data from this case, so it remains unknown whether this result is representative of a biological phenomenon or whether more complex constraints are required to capture the switch to Mode 2. In Mode 3, strain SAD’s efficiency is ∼5× lower than strain DAD’s due to its lack of fumarate reductase. Overall, efficiencies are higher during CF dechlorination than DCM mineralization, which is also reflected by experimentally determined yields (Table 1).

**Table 3.** Theoretical thermodynamic constraints for energy metabolism of both *Dehalobacter* models. Therm. max = maximum calculated using thermodynamics; FBA max = maximum calculated using flux balance analysis: NGAM = [optimized for] non-growth associated maintenance, Biomass = [optimized for] biomass. Energy transfer efficiency = (model-predicted ATP/*e^-^*)/(max ATP/*e^-^*).

| Strain | Growth mode<br>(donor/acceptor) | $\Delta G$<br>(kJ/e<sup>-</sup>) | ATP/e<sup>-</sup> ratio | | | | H<sup>+</sup>/e<sup>-</sup> ratio* | |
| --- | --- | --- | --- | --- | --- | --- | --- | --- |
|  |  |  | Therm.<br>max | FBA max |  | Energy<br>transfer<br>efficiency | Therm.<br>max | FBA<br>max |
|  |  |  |  | NGAM | Biomass |  |  |  |
| SAD | 1 (H<sub>2</sub>/CF) | -85 | 1.73 | 0.50 | 0.22 | 0.29 | 6.92 | 2.00 |
| DAD |  |  |  | 0.50 | 0.29 | 0.29 |  | 2.00 |
| SAD | 2 (DCM/CF) | -134 | 2.73 | 0.50 | 0.42 | 0.18 | 10.9 | 2.00 |
| DAD |  |  |  | 0.50 | 0.34 | 0.18 |  | 2.00 |
| SAD | 3 (DCM/H<sup>+</sup>) | -49 | 1.00 | 0.06 | 0.12 | 0.06 | 3.99 | 0.25 |
| DAD |  |  |  | 0.06 | 0.12 | 0.06 |  | 0.25 |

\* Assuming 4 protons are required per ATP synthesis

### Unexplained activity evidenced by Mode 3 redox imbalance

To further assess metabolic differences between these strains, flux balance analysis (FBA) was used to optimize each model, and solutions were explored under three metabolic modes. All FBA simulations are described and reported in Tables S9-S10, with summary of key fluxes in Table S11 (described in Supplemental Text S3). During Mode 3, optimization of each model using DCM as a sole electron donor and carbon source was initially infeasible due to stoichiometric redox constraints preventing ATP production (Figure 3), despite experimental growth of both strains under these conditions. To harness ATP synthase, one of two reactions must establish a proton motive force (Figure 3): a complex I-like oxidoreductase (FDXMQpp) or an energy conserving hydrogenase (HYDA_Ech). Computationally, neither translocation mechanisms can occur in the model during Mode 3, as described below.

**Figure 3.**
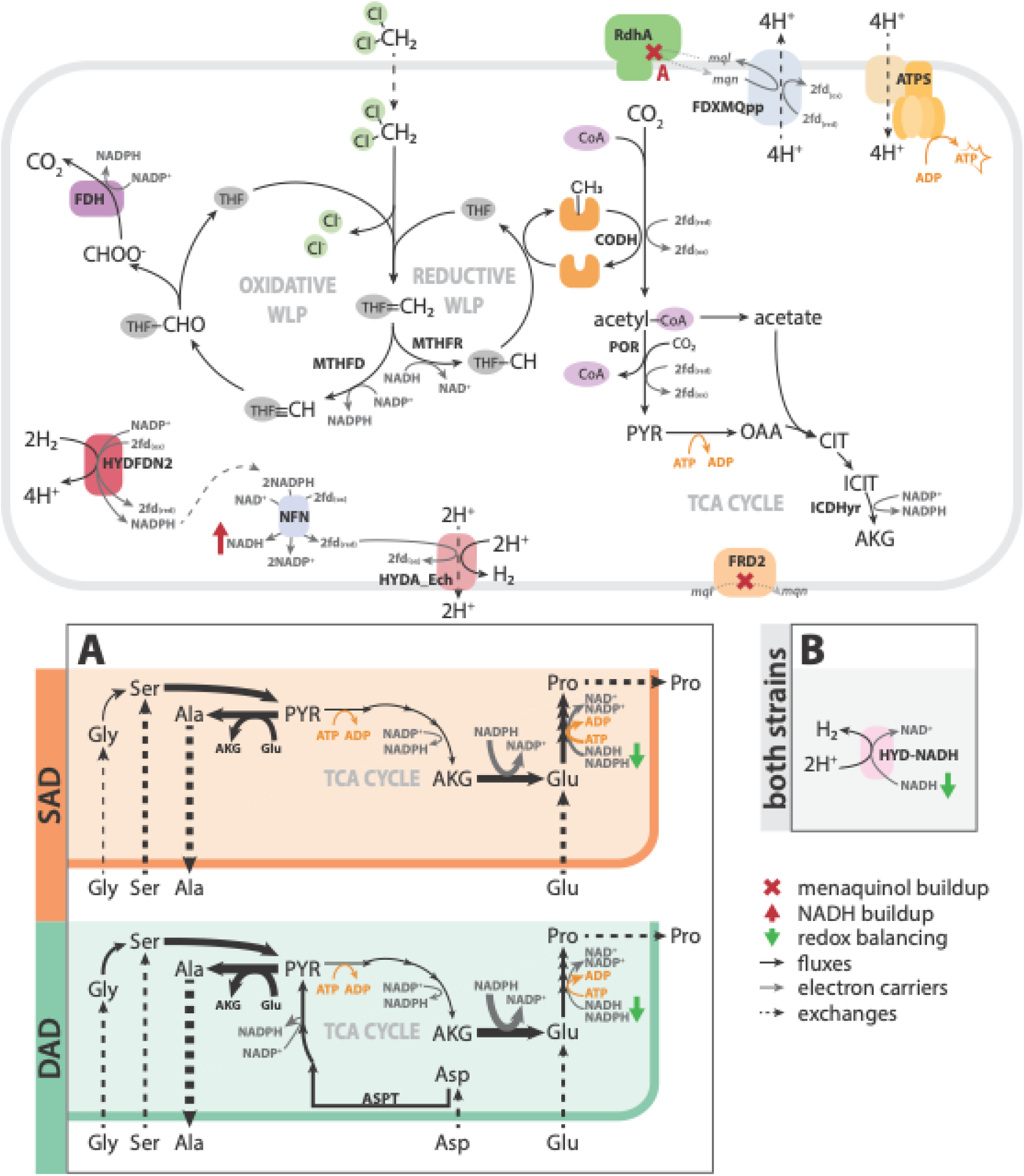
Redox imbalance during Mode 3 and strategies to resolve it (Panels A and B). Strategy A) amino acid supplementation to balance menaquinone pool in each strain. Strategy B) NADH-dependent hydrogenase. Metabolites are abbreviated as follows: PYR: pyruvate, ICIT: isocitrate, AKG: alpha-ketoglutarate, SUCC: succinate, FUM: fumarate, MAL: malate, THF: tetrahydrofolate, fdx: ferredoxin, mql: menaquinol, mqn: menaquinone. Enzymes are abbreviated as follows: MTHFD: methylene-THF dehydrogenase, MTHFR: methylene-THF reductase, CODH: carbon monoxide dehydrogenase/acetyl-CoA synthase, POR: pyruvate oxidoreductase, ICDHyr: isocitrate dehydrogenase, OORr: oxoglutarate oxidoreductase, FRD2: fumarate reductase, ME2: malic enzyme, NFN: electron-bifurcating transhydrogenase, HYDA_Ech/HYDFDN2/H2CYTBr/HYD-NADH: hydrogenases, FDXMQpp: complex I-like enzyme, RDase: reductive dehalogenase, ATPS: ATP synthase. WLP: Wood-Ljungdahl Pathway, TCA Cycle: Tricarboxylic acid cycle.

FDXMQpp requires an active menaquinol sink, which is limited to two possible reactions in the strain DAD model and only one in strain SAD. The RDase uses menaquinol to reduce CF to DCM in both strains, which is inherently inoperable without providing CF. In strain DAD, a putative fumarate reductase (FRD2) can use menaquinol to catalyze fumarate reduction to succinate, but strain SAD lacks this gene (Figure S1). Consequently, FDXMQpp cannot reduce menaquinone, and cannot function as a proton pump.

Alternatively, HYDA_Ech oxidizes ferredoxins to produce H_2_ and translocate protons. Despite the presence of ferredoxin recycling reactions in the model, NADH accumulation precludes a feasible solution. Ferredoxins can be reduced via two main reactions—a Group A3 [NiFe] hydrogenase (HYDFDN2) that reversibly bifurcates (or confurcates) electrons between two equivalents of H_2_ and one equivalent each of NAD(P)^+^ and oxidized ferredoxins, and a ferredoxin-dependent transhydrogenase (NFN) that bifurcates electrons from two equivalents of NADPH to reduce oxidized ferredoxins and NAD^+^. Due to this strict stoichiometry and the absence of an NADH sink, especially in the absence of FDXMQpp, a feasible solution is unattainable during Mode 3.

Three strategies resolved the redox imbalance imposed by DCM assimilation. First, supplementation of additional carbon sources like amino acids can absorb some of the accumulated reducing equivalents (Strategy A, Figure 3A). Alternatively, a hydrogenase or redox protein may produce H_2_ using the accumulated of NAD(P)H directly (Strategy B, Figure 3B), perhaps in conjugation with amino acid supplementation (Strategy C).

### Strategy A: Carbon source supplementation reconciles redox imbalance

The models were first supplemented with 5 mmol gdw^-1^ d^-1^ of each amino acid to reconcile the redox imbalance created during DCM assimilation and support biomass formation via conversion to pyruvate (Figure 3A). This supplementation resulted in feasible growth solutions in both strains (Figure S3, Table S10). To produce biomass from 10 mmol gdw^-1^ d^-1^ DCM, strain SAD required at least 5 mmol gdw^-1^ d^-1^ of glutamate and serine as well as smaller amounts of glycine, glutamine, and aspartate (Table S11). Strain DAD could also incorporate 5 mmol gdw^-1^ d^-1^ aspartate due to its aspartate transferase gene (ASPT), which doubled the growth rate in this simulation (Table S11). This increased substrate flexibility allowed biomass production feasibility with lower uptake fluxes of amino acids overall compared to strain SAD (3 mmol) (Table S11, Figure S3B).

### Strategy B: Hydrogen as an alternative NAD(P)H sink

As an alternative to supplementation with external carbon sources, the wealth of hydrogenases in each *Dehalobacter* genome provides a putative sink for reducing equivalents. When NADH-dependent hydrogen production (HYD-NADH) is added to the *Dehalobacter* models, both can produce biomass from DCM alone, without amino acid supplementation, indicating a successful NADH sink (Figure S4, Table S10). Growth rates under this condition are the same for each strain (0.007 d^-1^; Table S11), and both *Dehalobacter* genomes encode three homologous Group A3 [FeFe] hydrogenases: Hyd-3, Hyd-4, and Hyd-6 (Table S6). The three Group A3 [FeFe] hydrogenase β-subunits exhibit two different conserved domain patterns (Figure 4). The Hyd-3 β-subunit corresponds with that of electron-bifurcating hydrogenases, though it possesses an extra NAD/FAD-binding domain similarly to that of the *Sporomusa*-type NFN bifurcating oxidoreductase (25). The β-subunits of Hyd-4 and Hyd-6 align with the non-bifurcating hydrogenase of *S. wolfei,* which only has one flavin-binding domain and one FeS cluster (Figure 4) and may indicate their NAD(P)H-dependent activity.

**Figure 4.**
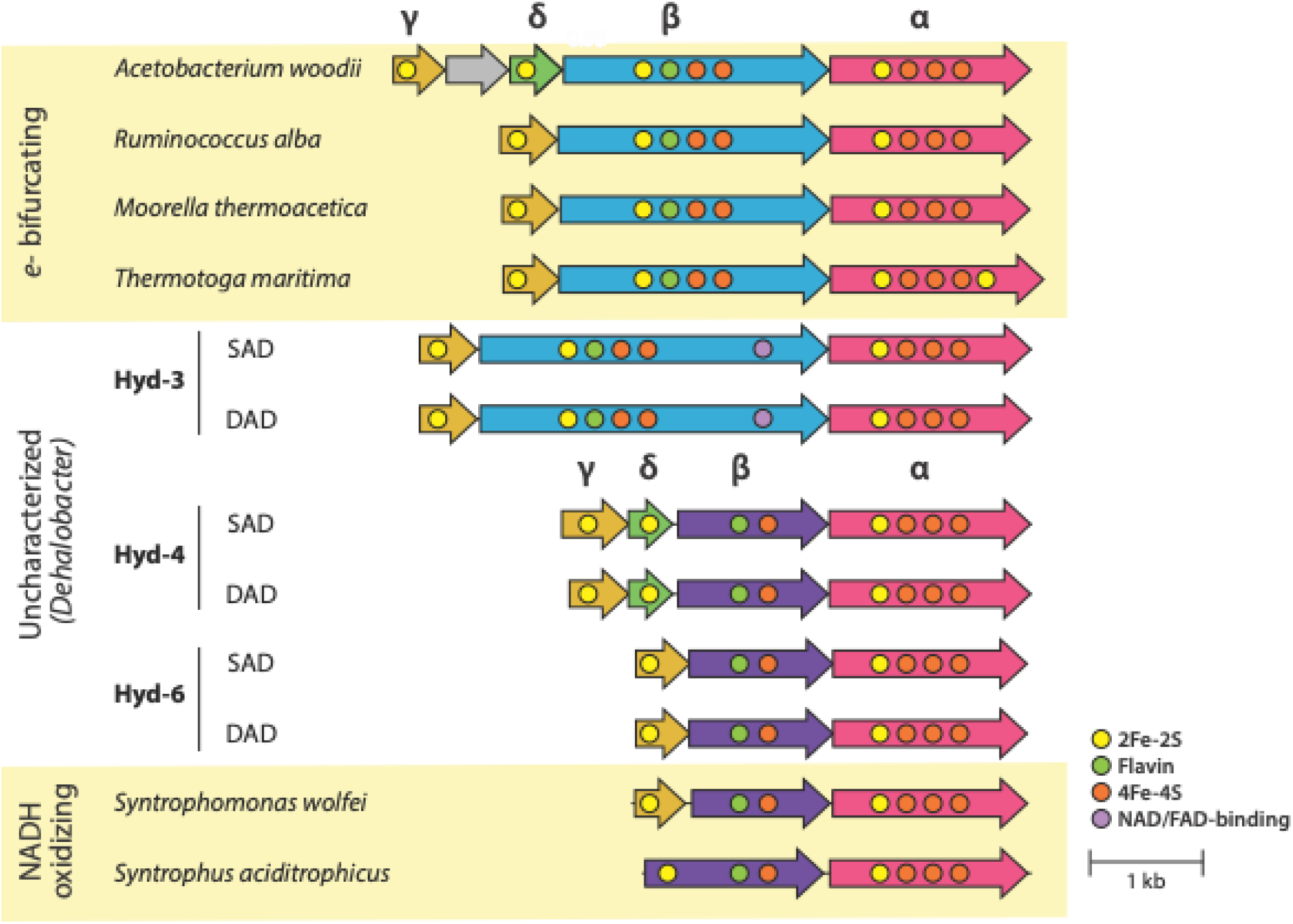
Group A3 [Fe-Fe] hydrogenases in *Dehalobacter* strains SAD and DAD and their predicted conserved motifs compared to characterized *e^-^* bifurcating and NADH-oxidizing hydrogenases (26). Subunits are colored by homology, as clustered using clinker (27).

### Strategy C (a combined approach) most accurately describes *Dehalobacter* growth

Experimental yields from this study and from the literature were compared to the FBA-predicted yields for each model (Table 4). During DCM mineralization, all predicted Mode 3 yields are within the same order of magnitude as experimental yields (Table 4). Strategies A and C are slightly higher, due to the supplement of amino acids, especially in strain DAD. Hydrogen export has been measured in SC05 (16), but is only predicted by the model when using an NADH-dependent hydrogenase (Strategies B and C, Table S11), nominating this strategy as the most phenotypically accurate. Overall, a combined strategy using an NADH-dependent hydrogenase with additional carbon source supplementation (Strategy C) may occur in the SC05-UT and DCME cultures, to explain experimental hydrogen production and the strains’ differential success under DCM mineralization conditions in DCME, though factors external to reaction stoichiometry may also drive community composition in this ecosystem, as discussed further below.

**Table 4.** Comparison of yields for each *Dehalobacter* strain during each growth mode, as determined experimentally and through flux balance optimization (FBA). ND = not determined; NF = not feasible. Solutions that predict hydrogen export are bolded.

| | | | Experimental<br>($\times 10^{12}$ cells/ eeq donor) | | FBA Solution<br>($\times 10^{12}$ cells/ eeq donor) | | | |
| --- | --- | --- | --- | --- | --- | --- | --- | --- |
| Strain | Mode |  | This study | Literature<sup>1</sup> | None | A | B | C |
|  |  |  |  |  |  | (*amino acids*)<sup>2</sup> | (*HYD-NADH*) | (*HYD-NADH + amino acids*)<sup>3</sup> |
| SAD | 1 | H<sub>2</sub>/CF | 13.1 ± 4.0 | 24.6 ± 1.5 | 3.3 | 5.6 | 9.8 | 18.8 |
|  | 2 | DCM/CF | ND | ND | NF | 0.1 | **8.1** | **11.7** |
| | 3 | DCM/H<sup>+</sup> | 6.8 | 5.3 ± 2.8 | NF | 3.3 | **1.3** | **2.7** |
| DAD | 1 | H<sub>2</sub>/CF | 37.1 ± 7.8 | 12.3 ± 6.5 | 6.1 | 14.2 | 15.1 | 29.3 |
|  | 2 | DCM/CF | ND | ND | 6.1 | 14.2 | **9.8** | **20.7** |
| | 3 | DCM/H<sup>+</sup> | 9.4 ± 3.0 | 8.0 ± 1.0 | NF | 6.2 | **1.4** | **3.2** |

<sup>1</sup>Taken from Bulka *et al.* (15), assuming all *Dehalobacter* growth in SC05-UT is strain SAD and all *Dehalobacter* growth in DCME is strain DAD, as determined in Bulka *et al.* (5).
<sup>2</sup>Amino acid fluxes used are 1 mmol gdw<sup>-1</sup> d<sup>-1</sup> for modes 1 and 2, and 5 mmol gdw<sup>-1</sup> d<sup>-1</sup> for mode 3.
<sup>3</sup>Amino acid fluxes used are 1 mmol gdw<sup>-1</sup> d<sup>-1</sup> for all modes.

### Intracellular electron transfer mediates self-feeding during CF dechlorination

SC05-UT is fed CF alone without exogenous H_2_ (Mode 2), during which electron cycling could occur two ways. The first is transmembrane hydrogen cycling, wherein a [NiFe] Group 1a Hup-type periplasmic uptake hydrogenase (H2CYTBr) uses periplasmic H_2_ to dechlorinate CF to DCM, which is then mineralized to produce cytosolic H_2_. After diffusion to the periplasm, this H_2_ is re-oxidized by H2CYTBr for further dechlorination of CF (Figure S4). FBA predicts transmembrane hydrogen cycling as the main electron transfer mechanism in strain SAD (Figure 5A); each of these simulations predict H_2_ export.

**Figure 5.**
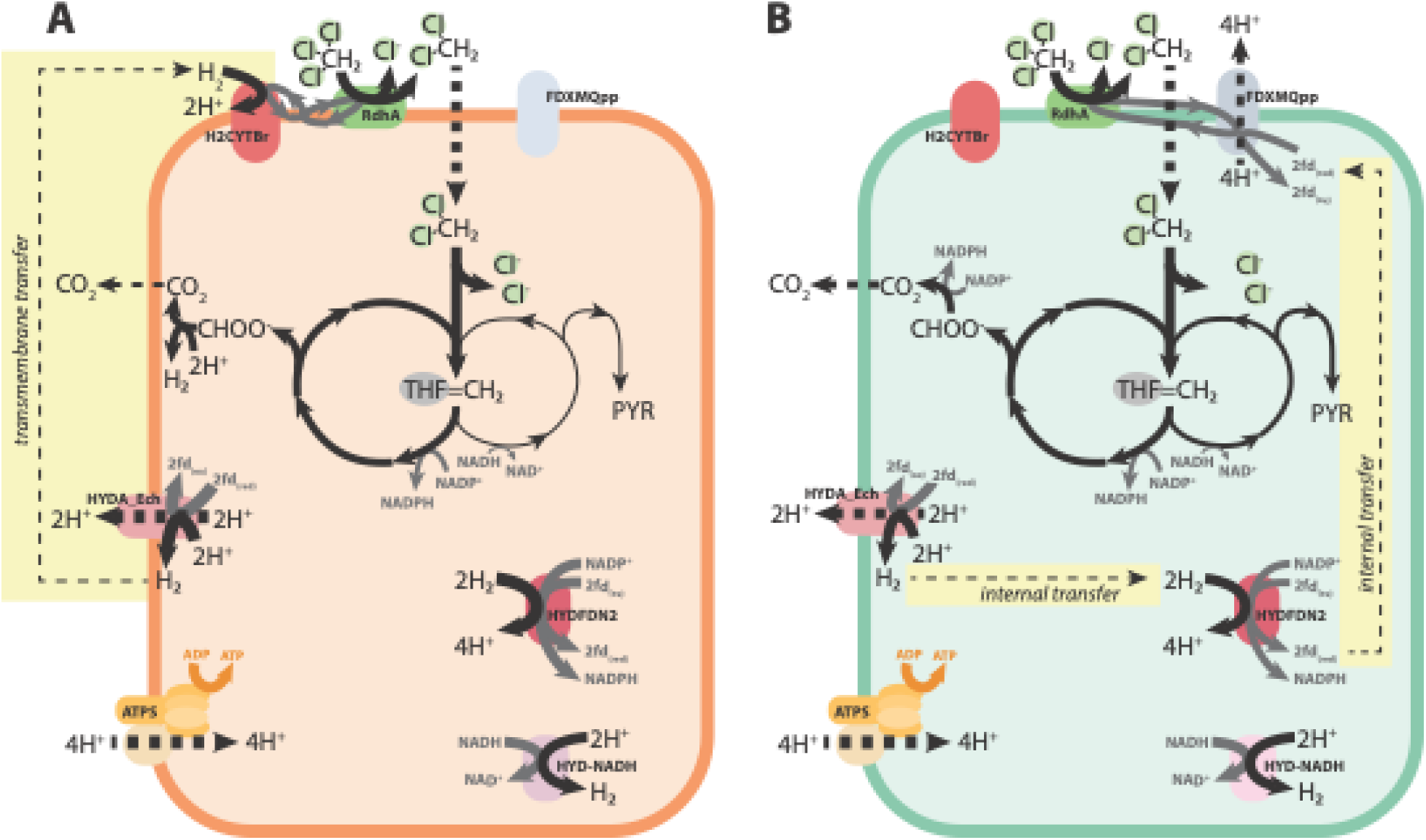
FBA-predicted electron transfer in *Dehalobacter* strains, including A) transmembrane transfer, and B) internal transfer. Arrows are weighted by relative flux. Metabolites and enzymes are abbreviated as follows: THF: tetrahydrofolate, fdx: ferredoxin, HYDA_Ech: Ech-type hydrogenase, HYDFDN: electron- bifurcating hydrogenase, H2CYTBr: cytochrome B-reducing hydrogenase, HYD-NADH: NADH-dependent hydrogenase, FDXMQpp: complex I-like enzyme, RDase: reductive dehalogenase, ATPS: ATP synthase.

In many simulations, HYDFDN is the main uptake hydrogenase. In this process, the DCM mineralization phase leads to production of H_2_ by HYDA_Ech or HYD-NADH, which accumulates until the second phase, wherein the hydrogen pool is recycled by HYDFDN to donate *e-* back to CF (Figure 5B). In these instances, H_2_ can be shuttled through a recycling loop between HYDA_Ech (H_2_ producing), HYD-NADH (H_2_ producing), and HYDFDN (H_2_ consuming) (Figure 5B). Notably, FBA models do not inherently consider temporal concentration dynamics (i.e. the consumption of hydrogen by HYDFDN may first require the accumulation of hydrogen in the system), so simulations were also performed with H_2_ consumption by HYDFDN inhibited (Table S10, solutions 17-22). In these simulations, transmembrane hydrogen cycling only occurs in strain SAD (Strategy A: with amino acids and no NADH-dependent hydrogenase); all other simulations report intracellular electron transfer using reduced ferredoxins that were produced from DCM assimilation to reduce menaquinone (FDXMQpp) and subsequently dechlorinate CF. Support for these mechanisms is discussed in later sections.

## Discussion

### Amino acid exchange in SC05

In a complex community like SC05, metabolites are exchanged between diverse organisms. For example, several amino acid-recycling microbes (the Dehalobacteriia class and Spirochaetes) are prominent in DCME (15). The previously modeled *Ca.* D. alkaniphilus CF relied on many metabolites produced by microbes in its community to resolve a similar redox imbalance (6), and supplemental amino acids, yeast extract, or spent media have also aided isolation and enrichment of many *Dehalobacter* strains (6, 7, 9, 22). The heavier reliance of strain SAD on additional supplements for DCM mineralization may contribute to the prominence of strain DAD in DCME. While strain SAD is evidently competitive in CF-fed conditions, strain DAD can produce more biomass from DCM alone with less support from amino acid–producing microorganisms because of the increased ability to harness its incomplete TCA cycle.

### Non-canonical hydrogenases in *Dehalobacter*

In each *Dehalobacter* genome are three Group A3 [FeFe] hydrogenases (Table S6). Most Group A3 [FeFe] hydrogenases bifurcate electrons, as was first demonstrated in *Thermotoga maritima*, which uses equimolar amounts of NADH and reduced ferredoxin to produce hydrogen (28). Bifurcating [FeFe] hydrogenases have also been studied in *Acetobacterium woodii* (29), *Moorella thermoacetica* (30), and *Ruminococcus albus* (31). Under low partial pressures (<10 Pa), however, the redox potential of hydrogen is about −260 mV, rendering hydrogen production from NADH (E′ = –320 mV) thermodynamically feasible even without confurcation (32).

Precedent for non-canonical, non-bifurcating behaviour exists for [FeFe] hydrogenases in several species. In *Syntrophomonas wolfei*, a Group A3 [FeFe] hydrogenase was shown experimentally to re-oxidize NADH to produce H_2_ independently of ferredoxins (26). *Syntrophus aciditrophicus* (33) and *Desulfovibrio fructosivorans* (34) also harbor characterized non-bifurcating NAD(P)H-dependent Group A3 [FeFe] hydrogenases, though the latter has only been characterized in the NADP-reducing direction. An [FeFe] hydrogenase from *Caldanaerobacter subterranus* (formerly *Thermoanaerobacter tengcongensis*) was also originally reported to produce hydrogen from NADH alone (35), but further investigation revealed its dependence on an additional electron acceptor (i.e. ferredoxin), which had been inadvertently supplied as titanium citrate in the original assays (36). More hydrogen was produced by the *S. wolfei* and *S. aciditrophicus* hydrogenases when supplied a higher NADH/NAD^+^ ratio, suggesting NADH accumulation facilitates more efficient hydrogen evolution (26, 33); however, hydrogen production by the *S. aciditrophicus* hydrogenase has been reported at an NADH/NAD^+^ ratio as low as 0.2 (33).

The β-subunits of these NAD(P)H-dependent [FeFe] hydrogenases and their electron-bifurcating counterparts comprise different conserved domains, corresponding with their different electron transfer reactions (26, 33). In NAD(P)H-dependent hydrogenases, the β-subunits are smaller and lack two FeS clusters compared to that of their electron-bifurcating counterparts, which is also consistent with NAD(P)H-dependent and electron-bifurcating formate dehydrogenases (26, 33). Similar β-subunits are seen in *Dehalobacter* Hyd-4 and Hyd-6 (Figure 4), which suggests that these hydrogenases also oxidize NADH to produce hydrogen—especially if NADH accumulates to high concentrations in the cell—and proposes a mechanistic explanation for consumption of excess electron equivalents in *Dehalobacter*.

### Model-predicted hydrogen cycling mirrors other domains of life

During CF dechlorination and DCM mineralization, both *Dehalobacter* models predict proton translocation through its Ech-type hydrogenase (HYDA_Ech) or the complex I-like oxidoreductase (FDXMQpp) (Figure 4). Both necessitate a source of reduced ferredoxins, whose production is predicted via a hydrogenase cascade (Figure 6). The model-predicted transmembrane hydrogen cycling (Figure 4A) mirrors the hydrogen cycling model that was first proposed to explain transient H_2_ observed in batch culture of *Desulfovibrio* spp. (Figure 6B) (37). In this model, a cytoplasmic hydrogenase converts electrons and protons generated during lactate fermentation to H_2_, which subsequently diffuses to the periplasm. A periplasmic hydrogenase re-oxidizes H_2_, establishing a proton gradient that drives ATP production. Simultaneously, electrons flow through several membrane-bound electron transfer complexes to ultimately reduce sulfate. This model has since expanded to include additional hydrogenases and H_2_- independent electron transfer to the electron transport chain via the complex I-like oxidoreductase or unknown electron transfer complexes (like Figure 4B) (34, 38–41).

**Figure 6.**
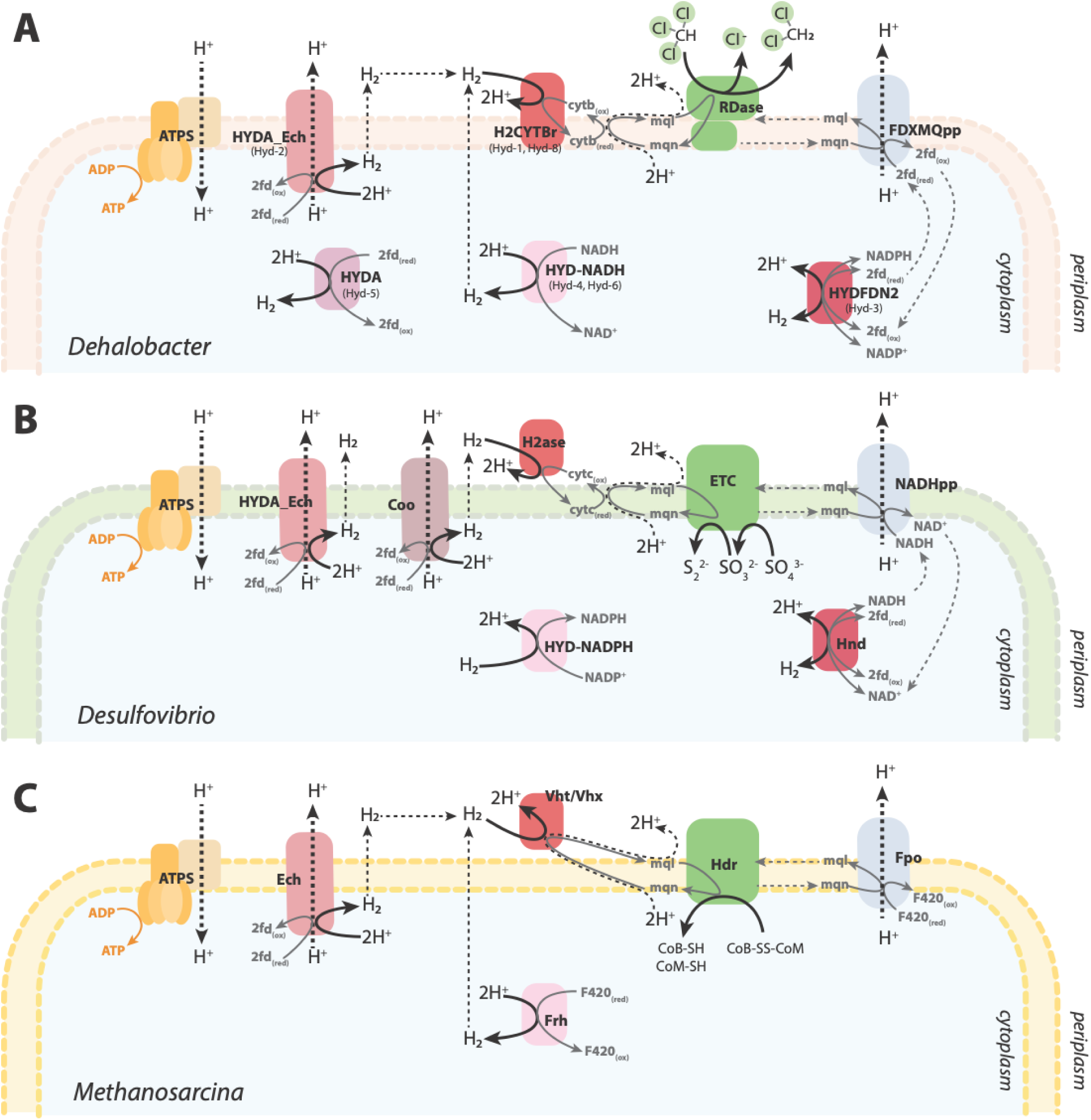
Proposed schematic for intracellular hydrogen cycling (without fluxes or stoichiometry) in **A**) *Dehalobacter* compared to characterized strains: **B**) *Desulfovibrio* (48) and **C**) *Methanosarcina* (49, 50). Metabolites are abbreviated as follows: fdx: ferredoxin, mql: menaquinol, mqn: menaquinone, cytb: cytochrome b, CoB: coenzyme B, CoM: coenzyme M. Enzymes are abbreviated as follows: H2ase: hydrogenase. HYDA_Ech: Ech-type hydrogenase, HYDFDN: electron-bifurcating hydrogenase, H2CYTBr: cyt B–reducing hydrogenase, HYD-NADH: NADH-dependent hydrogenase, NADHpp/FDXMQpp: complex I-like enzyme, RDase: reductive dehalogenase, ATPS: ATP synthase, ETC: electron transfer proteins.

Similarly, in *Methanosarcina barkeri*, the role of H_2_ cycling in energy conservation has been determined experimentally (Figure 6C) (42, 43). Here, H_2_ is produced by Ech and Frh (a F420-oxidizing cytoplasmic hydrogenase), which diffuses to the periplasm and is re-oxidized by Vht (a quinone-reducing periplasmic hydrogenase) to form a proton gradient harnessed by ATP synthase. An analogous system also exists in *Methanosarcina mazei* (44), and resembles the *Dehalobacter* transmembrane H_2_ cycling. The common occurrence of both cytoplasmic and periplasmic hydrogenases among diverse microorganisms suggests that this mechanism is widespread in nature, especially among anaerobes (42).

Without a periplasmic hydrogenase, like in fermenting *Pyrococcus* and *Thermococcus* strains, cytosolic H_2_ is produced by Ech and later re-oxidized by soluble [NiFe] hydrogenases to regenerate redox cofactors from what is otherwise a waste product (45, 46). This temporal intracellular hydrogen recycling also occurs in *Solidisulfovibrio fructosivorans* (formerly *Desulfovibrio fructosivorans*) during fermentation, where a bidirectional [FeFe] electron bifurcating hydrogenase (Hnd) reversibly produces hydrogen as an electron sink when grown under fermentative conditions and re-consumes it under respiratory conditions when an alternative electron acceptor is available (47). This mirrors simulations of sequential DCM mineralization (Mode 3) and subsequent CF reduction upon refeeding (Mode 1) by *Dehalobacter*.

### Other possibilities for redox balancing and limitations of metabolic modelling

The architecture of the *Dehalobacter* electron transport chain is not fully elucidated. Though menaquinone has been crystalized between the RdhA and RdhB subunits of a *Dehalobacter* RDase (51), the role of the RdhC subunit is still unknown. Due to its structural similarity to subunits of the Nuo NADH:ubiquinone oxidoreductase (Complex I), and its loose association with the other RDase subunits and ETC-related hydrogenases, it has been posited as an electron-transfer protein (4). This subunit may be able to accept electrons from intermediates like NAD(P)H or ferredoxins more directly, which would affect the redox balance of the cell, but further studies must be performed to further characterize this subunit’s function.

Many factors may impact the metabolism of each strain, including differential resilience to toxicity (CF especially) or membrane constraints (considering membrane-bound respiratory machinery), as well as enzyme rates and specificity. There are five single nucleotide variations (SNVs) between the homologous RDases in these two strains (14), which may have an impact on CF binding (K_s_) or dechlorination rate (µ_max_). Heterologous expression of the expressed strain DAD RDase and subsequent enzyme kinetics must be performed to determine if these SNVs play a role in the *Dehalobacter* population dynamics in SC05. Enzyme assays to compare activity of RDases and Mec cassettes in each strain may shed more light on these dynamics, as might microscopy of the different strains to compare their surface area/volume ratios.

### Organohalide syntrophy vs. competition

Though FBA simulations are helpful for determining the metabolic capability of each strain alone, in the SC05-UT and DCME subcultures, both strains have co-existed for many years. These strains were initially expected to interact symbiotically through intercellular compartmentalization, where labor is divided between strains such that one specializes in CF dechlorination, producing DCM for the other strain, which performs DCM mineralization and produces H_2_ for the CF-dechlorinator. Sequential steps of other pollutant biotransformation pathways that produce inhibitory intermediates are often compartmentalized within different cells across the population (52–54). This scheme reduces metabolic burden, maximizes the community growth rate, and may theoretically circumvent buildup of toxic intermediates (55). Experimentally, however, some DCM accumulates during CF dechlorination in SC05, and enzymes specific to both steps in the pathway are expressed by both *Dehalobacter* strains, regardless of steps are occurring (i.e. the CF-specific RDase is still expressed after extended enrichment on DCM), thus this scheme does not accurately describe the population dynamics of SC05.

An alternate scheme poses these strains as competitors for CF and DCM, with internal cycling of electrons or a shared pool of H_2_ as an electron donor to continually reduce CF completely to carbon dioxide. Complete use of one substrate by a single organism theoretically maximizes yield of the community, reducing the need for redundant cell maintenance expenditures (55). The need to avoid buildup of toxic intermediates may not play a large role in these organisms, since CF dechlorination occurs extracellularly in the periplasm, allowing the cell to control import of DCM for mineralization intracellularly. Furthermore, metabolic modelling suggests that higher yields are feasible where each strain is consuming both CF and DCM (Mode 2), as predicted yields for DCM mineralization alone (Mode 3) are low (Table 4). Experimentally, each strain is capable of both CF and DCM utilization, yet there is some evidence for tandem degradation in SC05-UT replicate 2 (Figure 1). In this replicate, strain DAD did not grow while CF was converted to DCM, but suddenly flourished as DCM was degraded. This suggests organohalide syntrophy between the two strains is also feasible, likely dependent on the environmental conditions in the culture.

### Role of methanogens in SC05

The feasibility of DCM mineralization is dependent on environmental hydrogen concentration, likely due to enzyme inhibition of hydrogen evolving hydrogenases (56–58). Modelling of this relationship is expanded upon in Supplemental Text S3.2. Previously studied DCM mineralizers require a syntrophic partnership with hydrogen consumers—typically methanogens or acetogens—to keep the hydrogen concentration low enough to allow DCM degradation to proceed (56). These relationships are also certainly at play in SC05-UT and DCME; DCM mineralization coincides with methane production, suggesting hydrogen consumption by methanogens reverses inhibition of the hydrogenases that are required for DCM mineralization (Figure 7B).

**Figure 7.**
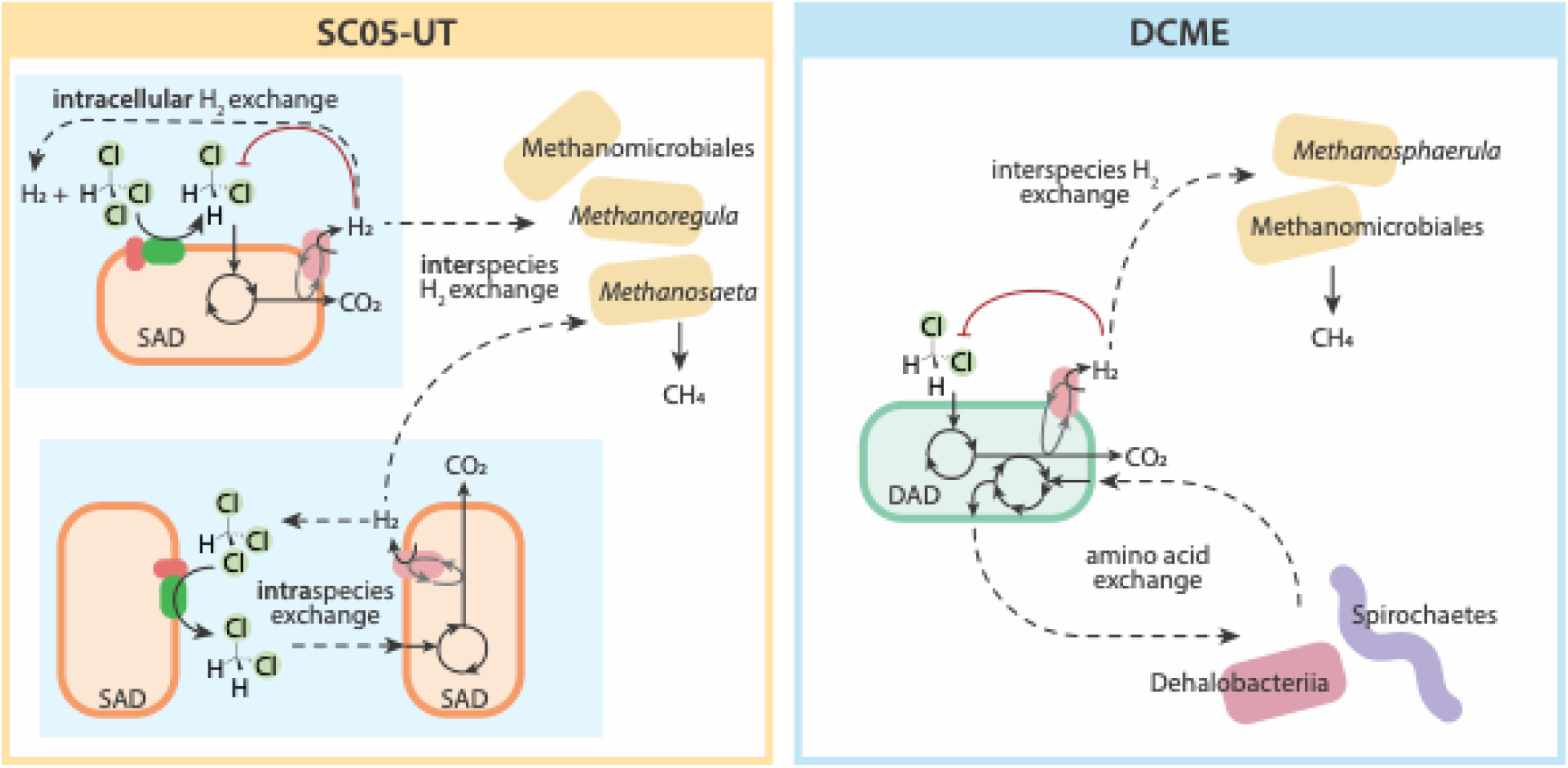
Proposed model of redox imbalance resolution of A) *Dehalobacter restrictus* SAD in SC05-UT and B) *Ca.* Dehalobacter alkaniphilus DAD in DCME, considering all data. Metabolite exchanges are depicted with dashed arrows; inhibition is depicted in red.

In the case of these two *Dehalobacter* strains, in Mode 2, hydrogen can be consumed as an electron donor to dechlorinate CF, maintaining a low hydrogen concentration for DCM mineralization. Twice as much hydrogen is produced from DCM mineralization as is consumed by CF dechlorination to DCM (15), which necessitates additional hydrogen consumption by methanogens. Methanogenesis becomes unfavorable at higher concentrations of hydrogen than dechlorination (Figure S7), which may provide an ecological advantage for a single multifunctional *Dehalobacter* strain able to consume both compounds, rather than requiring an interspecies exchange with a methanogenic organism.

### Significance for bioremediation

Due to lack of microbial isolates, the metabolic mechanisms of key microbes in the CF and DCM–degrading SC05 culture has remained elusive despite its wide use in bioaugmentation. By closing two distinct *Dehalobacter* genomes, and reconstructing metabolic models of each, we can better understand possible mechanisms of energy and biomass production during the multiple growth cases we observe experimentally. Moreover, studying the redox limitations of the pathways in each strain provides insight into caring for the cultures, and explanation as to its many possible growth phenotypes.

More broadly, the differential ability of *Dehalobacter* strains SAD and DAD to produce biomass from DCM without CF reduction is a case study for the impact of the function of single gene (fumarate reductase) on a seemingly unrelated pathway (DCM assimilation). Despite having similar genomes and sharing genes for key enzymes, the inclusion of fumarate reductase allows more efficient redox balancing and more growth in strain DAD. These results have implications for *in situ* biomarker tracking, as detection of the *mec* cassette from strain SAD *in situ* would not necessarily portend efficient DCM remediation.

## Materials and Methods

### Model reconstruction and curation

Model construction is expanded upon in Supplemental Text S1. Genome-scale metabolic models representing *Dehalobacter* strains SAD (*i*OB638) and DAD (*i*OB649) were drafted from model *i*HH623, a previously curated *Dehalobacter* model (12, 13). The reactions and compounds in the template model were converted to be in accordance with the BiGG database. The biomass equation was not changed from previous model construction, and includes 50% protein (using the amino acid composition of *Bacillus subtilis* protein [142]), 10% RNA, 5% DNA, 5% phospholipids, 25% peptidoglycan, and 5% ash (12).

A comparison of metabolic genes in the genomes of *Dehalobacter* strains SAD and DAD is expanded upon in Supplemental Text S2. These genomes were assembled as previously described (18) and annotated with MetaErg (60). Metabolic pathways were reconstructed with Kegg Mapper (61) and validated against the genome annotations to curate the models. Genes with metabolic annotations not included in the initial draft reconstruction were reviewed for inclusion using PaperBLAST (62). RDase and transport reactions were updated to reflect experimentally determined electron donors and acceptors. All putative hydrogenases were classified with HYDdb (24), and reactions were adjusted accordingly. Added or changed reactions are summarized in Table S4. Models are summarized in Table 2.

### Thermodynamic constraints

Additional curation was performed to account for thermodynamic constraints for the three electron donor/acceptor pairs used by SC05 (Supplemental Text S1). Briefly, the ΔG^°^*’* of each reaction was calculated using Eq. 1, where ΔG^°^*’_a_*and ΔG^°^*’_d_* are determined from the energies of formation of each compound in each half reaction assuming standard conditions (63). The maximum ATP/*e^-^* ratio was determined with Eq. 2 assuming ΔG’*_P_* = 50 kJ/ mol ATP (where *n* is the number of electrons transferred in the reaction). The maximum H^+^/*e^-^* ratio—the number of protons translocated across the membrane per mol *e^-^* transferred from donor to acceptor—was calculated by Eq. 3, assuming 4 protons are required per ATP formed by ATP synthase (64). The proton translocation stoichiometry in the model was adjusted such that when non-growth associated maintenance (NGAM) is maximized during flux balance analysis (FBA), the ATP/*e^-^* and H^+^/*e^-^* ratios fall within the thermodynamic constraints. Energy transfer efficiencies were estimated by comparing theoretical and experimental values (Table S3).

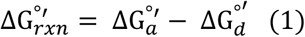

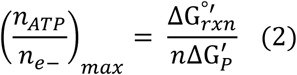

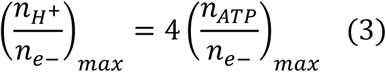

These parameters and experimental decay rates were used to calculate growth associated maintenance (GAM) and NGAM, which were ultimately set to 60 mmol ATP gdw^-1^ and 3.6 mmol ATP gdw^-1^ d^-1^, respectively (Supplemental Text S1.4). Substrate uptake rates were constrained to experimentally relevant values (10 mmol gdw^-1^ d^-1^) (63).

### Proteomic searching of *Dehalobacter* strains SAD and DAD

Reanalysis of pre-collected proteomic data (14, 65) was performed using the closed *Dehalobacter* strains SAD and DAD genomes (NCBI accessions: CP158031, CP148032 (18)). A description of protein extraction and mass spectrometry is described in our related work (14). All MS/MS samples were processed through X! Tandem (The GPM, thegpm.org; version X! Tandem Aspartate (2020.11.12.1)) for initial identification of peptides, and further analyzed through Scaffold (version Scaffold_5.3.0, Proteome Software Inc., Portland, OR), as described below.

X! Tandem was set up to search the custom database (6683 entries) assuming the digestion enzyme trypsin. X! Tandem was searched with a fragment ion mass tolerance of 0.40 Da and a parent ion tolerance of 20 PPM. Carbamidomethyl of cysteine and selenocysteine was specified in X! Tandem as a fixed modification. Cyclization of N-terminal glutamate and glutamine, ammonia-loss of the N-terminus, deamidation of asparagine and glutamine, and oxidation or dioxidation of methionine and tryptophan were specified in X! Tandem as variable modifications.

Scaffold was used to validate MS/MS based peptide and protein identifications. Peptide identifications were accepted if they could be established at greater than 95.0% probability by the Peptide Prophet algorithm (66) with Scaffold delta-mass correction (peptide FDR = 1.1%). Protein identifications were accepted if they could be established at greater than 90.0% probability and contained at least 1 identified peptide (protein FDR = 0.6%). Protein probabilities were assigned by the Protein Prophet algorithm (67). Proteins that contained similar peptides and could not be differentiated based on MS/MS analysis alone were grouped to satisfy the principles of parsimony.

### Flux balance analysis

Flux balance analysis (FBA) was performed using COBRApy (68). Constraints were applied to each model to render the simulations more physiologically relevant, considering several experimentally derived datasets: proteomic expression data (Dataset S2), metabolite profiling of the cultures, experimental growth rates, and dechlorination rates (15). Code and models can be found in the following GitHub repository: https://github.com/LMSE/Dehalobacter_modelling.

### Strain detection in each culture

Sub-transfers of SC05-UT and DCME were weekly set up as previously described (15). Briefly, three sub-transfers of SC05-UT and two sub-transfers of DCME were amended with 25 µmol CF (0.25 mM CF aqueous) and 10 µmol hydrogen (1 mL H_2_/CO_2_, 20:80, v/v) to jump-start degradation. The SC05- UT replicates were amended with another 50 µmol CF on day 4. Each bottle’s headspace was analysed by gas chromatography every 3-5 days, and 1 mL of culture was sampled for DNA extraction and qPCR.

Each strain of *Dehalobacter* was quantified using primers previously designed for a core gene in *Dehalobacter* with high sequence variability: flagellar basal-body rod protein, *flgC* (5). Using these primers, quantitative PCR (qPCR) was performed on DNA samples taken during previous time course experiments with the SC05-UT and DCME subcultures (15). The qPCR reaction mixtures were prepared in a UV-treated PCR cabinet (ESCO Technologies, Hatboro, PA), and contained 10 μL of 2× SsoFast SYBRGreen® (Bio-Rad, Hercules, CA), 0.25 μM of each *flgC* primer, and 2 μL of template DNA. The amplification program and analyses were conducted using a CFX96 Touch Real-Time PCR Detection System and the CFX Manager software (Bio-Rad). The qPCR method included an initial denaturation step at 98°C for 2 min, followed by 40 cycles of 5 s at 98°C and 10 s at 55°C, including a 2°C s^-1^ ramp between temperatures. Quantification was performed using 10-fold serial dilutions of PCR-produced standard DNA amplified from a larger region of the *flgC* gene. The number of copies was calculated assuming a 100% DNA extraction efficiency.

## Conflicts of interest

The authors declare that there are no conflicts of interest.

## Funding information

Funding for this work was provided by the Natural Science and Engineering Research Council (NSERC) though a Discovery Grant to E.A.E. and Doctoral Scholarships to O.B., as well as a Genome Canada Bioinformatics and Computational Biology (BCB project 285MPR) subgrant to E.A.E. and R.M. We also acknowledge funding from the Canada Research Chairs Program to R.M. and E.A.E. The funders had no role in study design, data collection, or interpretation.

